# Performance of phenomic selection in rice: effects of population size and genotype-environment interactions on predictive ability

**DOI:** 10.1101/2024.08.15.608050

**Authors:** H de Verdal, V. Segura, D. Pot, N. Salas, V. Garin, T. Rakotoson, L.M. Raboin, K. VomBrocke, J. Dusserre, S. Castro Pacheco, C. Grenier

## Abstract

Phenomic prediction (PP), a novel approach utilizing Near Infrared Spectroscopy (NIRS) data, offers an alternative to genomic prediction (GP) for breeding applications. In PP, a hyperspectral relationship matrix replaces the genomic relationship matrix, potentially capturing both additive and non-additive genetic effects. While PP boasts advantages in cost and throughput compared to GP, the factors influencing its accuracy remain unclear and need to be defined. This study investigated the impact of various factors, namely the training population size, the multi-environment information integration, and the incorporations of genotype x environment (GxE) effects, on PP compared to GP. We evaluated the prediction accuracies for several agronomically important traits (days to flowering, plant height, yield, harvest index, thousand-grain weight, and grain nitrogen content) in a rice diversity panel grown in four distinct environments. Training population size and GxE effects inclusion had minimal influence on PP accuracy. The key factor impacting the accuracy of PP was the number of environments included. Using data from a single environment, GP generally outperformed PP. However, with data from multiple environments, using genotypic random effect and relationship matrix per environment, PP achieved comparable accuracies to GP. Combining PP and GP information did not significantly improve predictions compared to the best model using a single source of information (e.g., average predictive ability of GP, PP, and combined GP and PP for grain yield were of 0.44, 0.42, and 0.44, respectively). Our findings suggest that PP can be as accurate as GP when all genotypes have at least one NIRS measurement, potentially offering significant advantages for rice breeding programs.

**Authors Summary:** This study explores the interest of phenomic selection within the context of rice breeding. Unlike genomic selection, phenomic selection utilizes near-infrared spectroscopic (NIRS) technology to predict genotype’s performance. The importance of this methodology lies in its capacity to reduce the costs and enhance the genetic gains of breeding programs, particularly in developing countries where genomic information is not always easily accessible (cost, availability, ease of use). Also, NIRS technology is often already available, even in resource-constrained breeding programs. By focusing the study on rice, a staple food for billions, our research aims to demonstrate the applicability of phenomic selection compared to genomic selection. By investigating the influence of various factors on phenomic prediction accuracy (training population size, incorporation of multiple environment information, consideration of genotype x environment effects in the prediction models), we are contributing to the optimization of this novel breeding method, which could potentially lead to significant improvements in agricultural productivity and food security.

## Introduction

Genomic selection (GS), a selection methodology based on genomic predicted values, has emerged as a powerful tool to enhance the efficiency of selection in plant and animal breeding (1). By leveraging genome-wide marker information, GS offers several advantages over traditional breeding methods. It can increase selection intensity and accuracy (2), leading to faster identification of superior genotypes. Additionally, GS can shorten the breeding cycle length (3,4), accelerating the development of varieties. The effectiveness of GS for predicting complex traits like grain yield in plants is well-established (5–8). Despite these advantages, GS adoption remains limited in a large share of breeding programs. One key limitation to broader GS adoption is the access and/or the cost associated with genotyping (1). Generating high-quality genotyping data for a large number of crosses at each selection cycle can be a significant technical and/or financial hurdle. While the genotyping costs have been steadily decreasing and the ease of genotyping has been generalized (9), this is mostly true in developed countries. In many breeding programs with limited resource (e.g. orphan crop, developing countries), access to efficient genotyping infrastructure remain complicated. Therefore, in such cases, the cost-effectiveness of GS deployment needs to be carefully evaluated.

Rincent et al. (10) proposed an innovative strategy aimed especially at optimizing breeding programs with limited access to genotyping facilities. This strategy involves the replacement of genomic information by hyperspectral information, measured for instance by near infrared reflectance spectrometry (NIRS), to capture genetic variance. Using this methodology, known as phenomic prediction (PP), the authors showed equivalent or even better PA than with genomic predictions (GP) for several important traits such as grain yield in wheat. NIRS is a high-throughput, low-cost and non-destructive method which is already frequently used in breeding programs to evaluate different biochemical traits for which wavelengths can serve as proxies, such as protein, starch, and oil content in grains or seeds (11). The reflectance spectra observed, likely arise from the complex interplay of various chemical bonds within the analyzed tissue, which depends on genetic effects but also on interaction between traits, epistatic interactions between specific loci and GxE effects (10).

In breeding programs, the aim is to deliver varieties adapted to growing conditions in target population of environments (TPE). Often, a variety specifically developed for one TPE may not perform as well elsewhere (12). This phenomenon, known as genotype-by-environment interaction (GxE), arises from the interplay between the plant’s genotype and the environment in which it is grown. A variety that exhibits superior performance in, for example, a research station might not deliver the same benefits in farmer’s fields, a problem which is likely to occur more frequently due to climate change (13,14). Integrating the knowledge acquired from GxE studies into breeding programs empowers breeders to develop varieties with improved adaptation to local environmental conditions. By incorporating GxE effects, breeders can move beyond performance in a single research station setting and identify varieties that consistently deliver desirable traits across diverse environments (15) or inversely, highly adapted to a specific TPE. The importance of GxE interactions is usually taken into account in the breeding scheme, through multi-environment trials (MET) (16). These METs consist of testing a given number of genotypes, generally new varieties of improved lines, in different sites to assess i) their stability across environments (or TPE) and ii) their potential in a specific TPE. One of the challenges associated with METs is the substantial cost resulting from evaluating genotypes across diverse locations (17), as these field trials are one of the most expensive components in breeding programs.

Furthermore, accurately predicting the performance of a genotype in a target environment where it has not yet been phenotyped remains a critical challenge (18). Development of prediction models integrating the GxE effects are useful in that sense (19–23), with an increase of predictive ability (PA) of these models compared to single-environment models. In this context, the use of genomic prediction (24–26) in which all the genotypes are not phenotyped in all the sites but predicted from genomic information is attractive, as it may reduce the phenotyping efforts and thus the global costs associated with MET (27,28).

Phenomic prediction with its potential to capture a portion of the genetic and GxE variations that contribute to spectra variability (10), offers the possibility to not only predicting traits directly related to the analyzed tissue, but also to predict genotype performance in contrasting environmental conditions. PS could be a good opportunity to improve breeding programs with reduced resources.

Another interesting aspect of hyperspectral measurement is that it can capture part of the GxE variance without additional environmental information which is not possible using genomic prediction (18,29). The NIRS profile of a single genotype can fluctuate between different environments (30). Recent literature has demonstrated the potential of phenomic prediction based on hyperspectral data. This approach has shown strong PA for various traits in several plant species, including soybean (31), maize (29), wheat (10,17,32), rye (33,34), sugarcane (35), coffee (36) and grapevine (37).

While PS has shown promise in various crops, its application in rice (*Oryza sativa*) breeding remains unexplored despite the importance of this crop worldwide. A comprehensive comparison of PS and GS is necessary to elucidate their relative advantages and limitations for rice breeding programs implementation (30). Therefore, the aim of the present study is i) to evaluate the potential of PS compared to GS in rice breeding using diverse cross-validation scenarios including between one and three environmental conditions and GxE interactions and ii) to investigate the influence of several factors on PS effectiveness, e.g., genetic architecture of traits, training population size and composition, and prediction models. These factors are well known to influence the accuracy of GP (2,38–41) and their impacts on PP remain to be evaluated (42).

The scenarios tested in the current study addressed three objectives: (a) predicting unknown genotypes in known environments, (b) predicting known genotypes in an unknown environment, (c) predicting known genotypes in known environments in which they are not observed (sparse testing scenario). All these scenarios were tested with a variable number of environments and different sets of genotypes included in the training populations. The dataset used originated from a diversity panel chosen to encompass the diversity used in the upland rice breeding program of FOFIFA and CIRAD institutes. This panel was grown in Madagascar in four contrasting environments corresponding to two nitrogen conditions, with high or low levels of nitrogen fertilizers, during two consecutive years (43).

## Results

### Phenotypic performances

Table 1 summarizes the descriptive statistics for the six investigated traits: days to flowering (DF), plant height (PH), harvest index (HI), thousand-grain weight (TGW), grain yield (GY) and grain nitrogen content (GNC). Number of days to flowering averaged 93.2 days with a standard deviation of 7.22 days. At maturity, average plant height was 102.5 ± 16.6 cm. Thousand grain weight averaged 28.2 ± 4.53 g and grain yield averaged 4,162 ± 1,490 kg.ha^-1^ with a coefficient of variation of 35.8%. The harvest index ranged from 0.04 to 0.64, and average GNC was 1.56 ± 0.18 %. Mixed linear models were conducted to partition the phenotypic variation for each measured trait into genotypic, genotype by year, genotype by nitrogen treatment, block and residual variances (Fig 1).

**Table 1.** Descriptive statistics of the phenotypic measurements from the four environments (2015-HN. 2015-LN. 2016-HN and 2016-LN) with mean, standard deviation (StdDcv). min, max, coefficient of variation (CV) and the broad-sense heritability (H^2^).

| Traits | Abbreviations | Mean | StdDev | Min | Max | CV | $H^2$ |
| --- | --- | --- | --- | --- | --- | --- | --- |
| Day to flowering (days) | DF | 93.2 | 7.22 | 70.0 | 116 | 7.76 | 0.85 |
| Plant height (cm) | PH | 102.5 | 16.6 | 49.3 | 162.2 | 16.2 | 0.94 |
| Thousand grain weight (g) | TGW | 28.2 | 4.53 | 14.9 | 44.0 | 16.1 | 0.94 |
| Grain yield ($\text{kg} \cdot \text{ha}^{-1}$) | GY | 4,162 | 1,490 | 227 | 8,871 | 35.8 | 0.58 |
| Harvest index | HI | 0.46 | 0.08 | 0.04 | 0.64 | 16.9 | 0.80 |
| Grain nitrogen content (%) | GNC | 1.56 | 0.18 | 0.09 | 1.19 | 29.4 | 0.60 |

**Fig 1.**
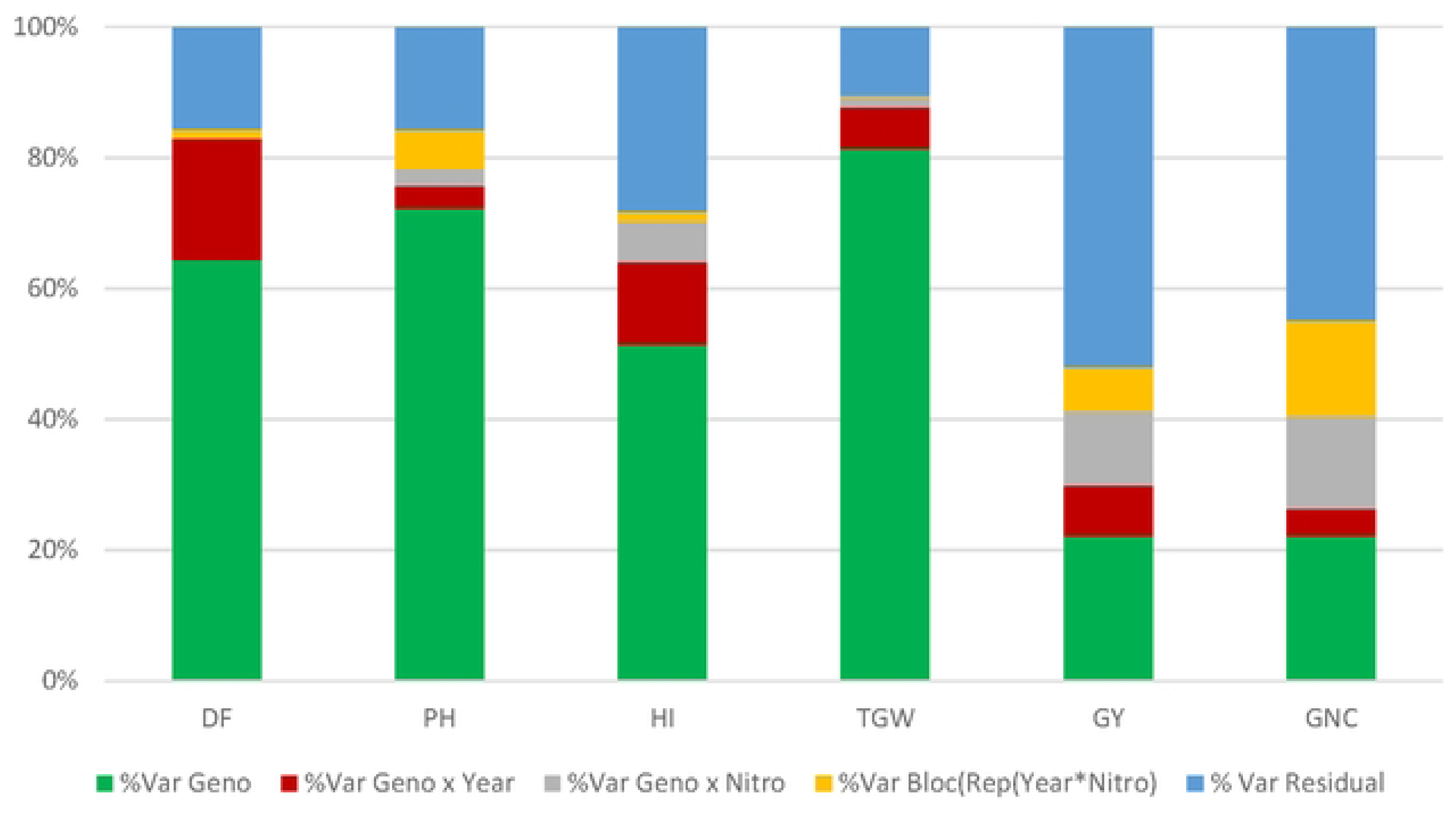
Variance decomposition for the six phenotypic traits’ across all the environments. Var Geno, genotypic variance; Var Geno x Year, variance of the genotype by year interaction; Var Geno x Nitro, variance of the genotype by nitrogen treatment interaction; Var Bloc(Rcp(Year x Nitro)), variance of the bloc nested in a replicate and a nitrogen treatment and year; and Var Residual, the residual variance. ‘DF. Day to flowering; PH. plant height; TGW. thousand grain weight; GY. grain yield; HI. harvest index; GNC. grain nitrogen content

Genotypic variance was moderate to high for all traits, ranging from 22.1% for GY and GNC to 81.4% for TGW. The interaction variances between genotypes and years, as well as between genotypes and nitrogen treatments, were low to moderate for all traits, ranging from 0% to 18.6%. Significant year effects were observed for all traits, and, in most cases except DF, exhibiting higher values in 2016 compared to 2015 (Table S1). Nitrogen fertilization in the field significantly impacted several traits except DF (Table S1). Plant height, GY and GNC all increased with high levels of nitrogen fertilizers (by 15.7 cm, 1,111 kg.ha^-1^ and 0.40%, respectively). Conversely, HN treatment significantly reduced TGW and HI by 0.51 g and 0.04, respectively. The block effect nested within replicate, year, and nitrogen treatment explained a relatively small portion of the total variance, ranging from 0.3% to 14.7%. Finally, broad sense heritability estimates were moderate to high, ranging from 0.58 for GY to 0.94 for PH and TGW (Table 1).

Table 2 presents the phenotypic correlations for each of the six traits based on their BLUEs measured across the four environmental conditions (2015-HN, 2015-LN, 2016-HN, and 2016-LN). High phenotypic correlations between environmental conditions, ranging from 0.56 to 0.93, were observed for DF, PH, TGW, and HI. Grain yield and GNC exhibited moderate correlations across environmental conditions, ranging from 0.22 to 0.61. These last two traits were those for which the interaction variances (genotype within a nitrogen treatment and genotype within a year variances) were high compared to the genetic variance.

**Table 2.** Pearson correlations between all pairs of environments for all the phenotypic traits’. Correlations were calculated between the BLUEs of the different environments, defined as a combination of year x nitrogen treatment (p-value < 0.0001)..

|  | 2015-HN<br>2015-LN | 2015-HN<br>2016-HN | 2015-HN<br>2016-LN | 2015-LN<br>2016-HN | 2015-LN<br>2016-LN | 2016-HN<br>2016-LN |
| --- | --- | --- | --- | --- | --- | --- |
| **DF** | 0,92 | 0,72 | 0,7 | 0,77 | 0,77 | 0,93 |
| **PH** | 0,87 | 0,87 | 0,83 | 0,81 | 0,82 | 0,87 |
| **TGW** | 0,93 | 0,87 | 0,84 | 0,88 | 0,87 | 0,92 |
| **GY** | 0,49 | 0,48 | 0,22 | 0,38 | 0,39 | 0,32 |
| **HI** | 0,8 | 0,66 | 0,56 | 0,67 | 0,67 | 0,68 |
| **GNC** | 0,43 | 0,61 | 0,24 | 0,39 | 0,42 | 0,42 |

<sup>1</sup>DF. Day to flowering; PH. plant height; TGW. thousand grain weight; GY. grain yield; HI. harvest index; GNC. grain nitrogen content

### Comparison of genomic, phenomic and multi-omics prediction models

#### Single environment models

While statistically different predictive abilities (PA) were observed across all cross-validation (CV) scenarios (S2 Table, S3 Table) within each environment, the magnitude of these differences was negligible. Furthermore, no consistent patterns emerged in PA differences related to either year or nitrogen treatment effects. Consequently, results from all environments were combined for further analysis (Fig 2).

**Fig 2.**
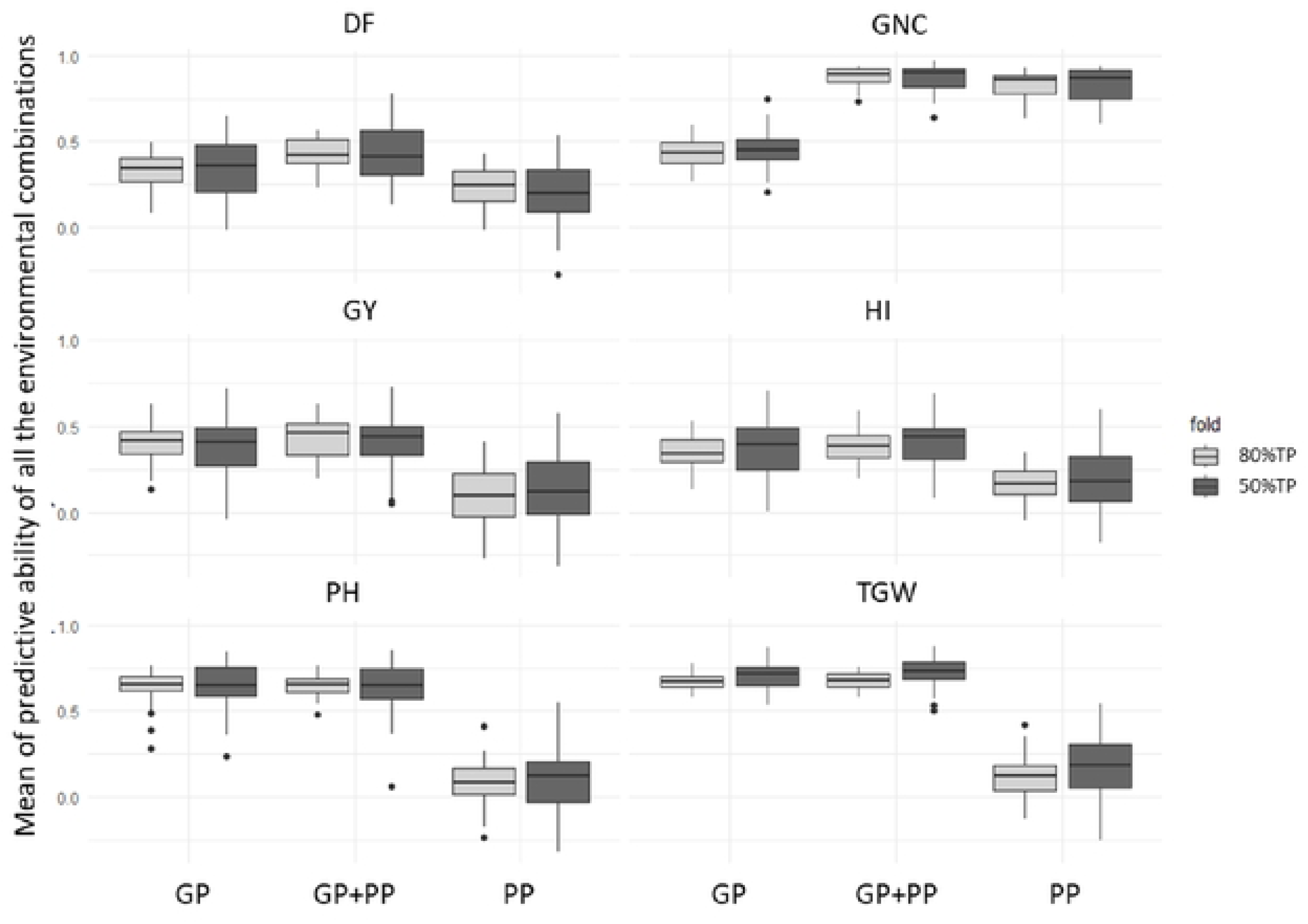
Means of the predictive ability for all the four environments when a single environment was considered in the training population. The dark grey boxes indicate the 2-fold cross-validation (50% of the population was in the TP and the other 50% in the VP), and the light grey boxes indicate the 5-fold cross-validation (80% of the population in the TP and the remaining 20% in the VP). The prediction models arc GP. genomic predictions; PP. phenomic predictions; GP+PP. combination of genomic and phenomic information for the predictions. The PAs arc averaged over the 4 scenarios tested (1 to 4. sec Table 1 for a detailed description of scenarios). DF. Day to flowering; PH. plant height; TGW. thousand grain weight; GY. grain yield; HI. harvest index; GNC. grain nitrogen content

For all traits except GNC, both GP and GP+PP delivered significantly higher PA compared to PP alone (Fig 2). Interestingly, while a larger training population improved PA (significant for PH, HI, TGW and GNC), this improvement appeared to be low (Fig 2). Finally, for GNC, GP+PP significantly improved PA compared to PP, which in turn performed significantly better than GP.

#### Multi-environments models

For scenarios involving two environments, all genotypes from the first environment were integrated into the TP as well as a variable proportion of genotypes from the second (targeted) environment. This proportion ranged from 0% (no genotype from the targeted environment were included in the TP) to 80%, corresponding to the number of folds used for cross-validation (Table 3). The remaining genotypes from the targeted environment formed the validation population (VP) and were subjected to prediction.

**Table 3.** Training and validation populations used for the different scenarios tested in the genomic and phenomic prediction approaches.

| Scenario | Number of environments<sup>†</sup> | Training population (TP) <sup>‡</sup> | Validation population (VP) <sup>‡</sup> |
| --- | --- | --- | --- |
| 1 | 1 | 2015-HN (50 or 80%) | 2015-HN (50 or 20%) |
| 2 |  | 2015-LN (50 or 80%) | 2015-LN (50 or 20%) |
| 3 |  | 2016-HN (50 or 80%) | 2016-HN (50 or 20%) |
| 4 |  | 2016-LN (50 or 80%) | 2016-LN (50 or 20%) |
| 5 | 2 | 2015-HN (100%) + 2015-LN (0. 10. 50 or 80%) | 2015-LN (100. 90. 50 or 20%) |
| 6 |  | 2015-LN (100%) + 2015-HN (0. 10. 50 or 80%) | 2015-HN (100. 90. 50 or 20%) |
| 7 |  | 2015-HN (100%) + 2016-HN (0. 10. 50 or 80%) | 2016-HN (100. 90. 50 or 20%) |
| 8 |  | 2016-HN (100%) + 2016-LN (0. 10. 50 or 80%) | 2016-LN (100. 90. 50 or 20%) |
| 9 |  | 2016-LN (100%) + 2016-HN (0. 10. 50 or 80%) | 2016-HN (100. 90. 50 or 20%) |
| 10 |  | 2015-LN (100%) + 2016-LN (0. 10. 50 or 80%) | 2016-LN (100. 90. 50 or 20%) |
| 11 | 3 | 2015-HN (100%) + 2015-LN (100%) + 2016-HN (0. 50 or 80%) | 2016-HN (100. 50 or 20%) |
| 12 |  | 2015-HN (100%) + 2015-LN (100%) + 2016-LN (0. 50 or 80%) | 2016-LN (100. 50 or 20%) |
| 13 |  | 2015-HN (100%) + 2016-LN (100%) + 2016-HN (0. 50 or 80%) | 2016-HN (100. 50 or 20%) |
| 14 |  | 2015-LN (100%) + 2016-HN (100%) + 2016-LN (0. 50 or 80%) | 2016-LN (100. 50 or 20%) |

<sup>†</sup> Refers to the total number of environments included in the scenario.
<sup>‡</sup> For scenarios 1 to 4, the k-fold CV was performed with k=5 (TP = 1-1/k = 80% and VP = 1/k = 20% of the population) or k=2 (TP = 50% and VP = 50% of the population). For scenarios 5 to 10 all the data from one environment were included in the TP as well as a proportion of 0. 10. 50 or 80% of the second environment. as a sparse testing model, the rest of the data of the second environment were accordingly allocated to the VP (100. 90. 50 or 20%). Finally for scenarios 11 to 14, all the data from two environments as well as three proportions (0. 50 or 80%) of the third environment were considered in the TP and the rest of the data of this third environment allocated to the VP.

Across all phenotypic traits and regardless of the relationship matrix employed (genomic - GP, phenomic - PP, or both combined – GP+PP), the inclusion of genotypes from the targeted environment in the TP significantly increased PA (S2 Table, S3 Table). Furthermore, a tendency of PA improvement was observed with the increase of the percentage of genotypes included in the targeted environment for the TP (Fig 3). Moreover, the models integrating GxE effect (MDs model) tended to deliver comparable or improved PA compared to models without GxE (MM model), regardless of the other effects tested (Fig 3) for all traits except DF and PH. While significant, these differences stayed low.

**Fig 3.**
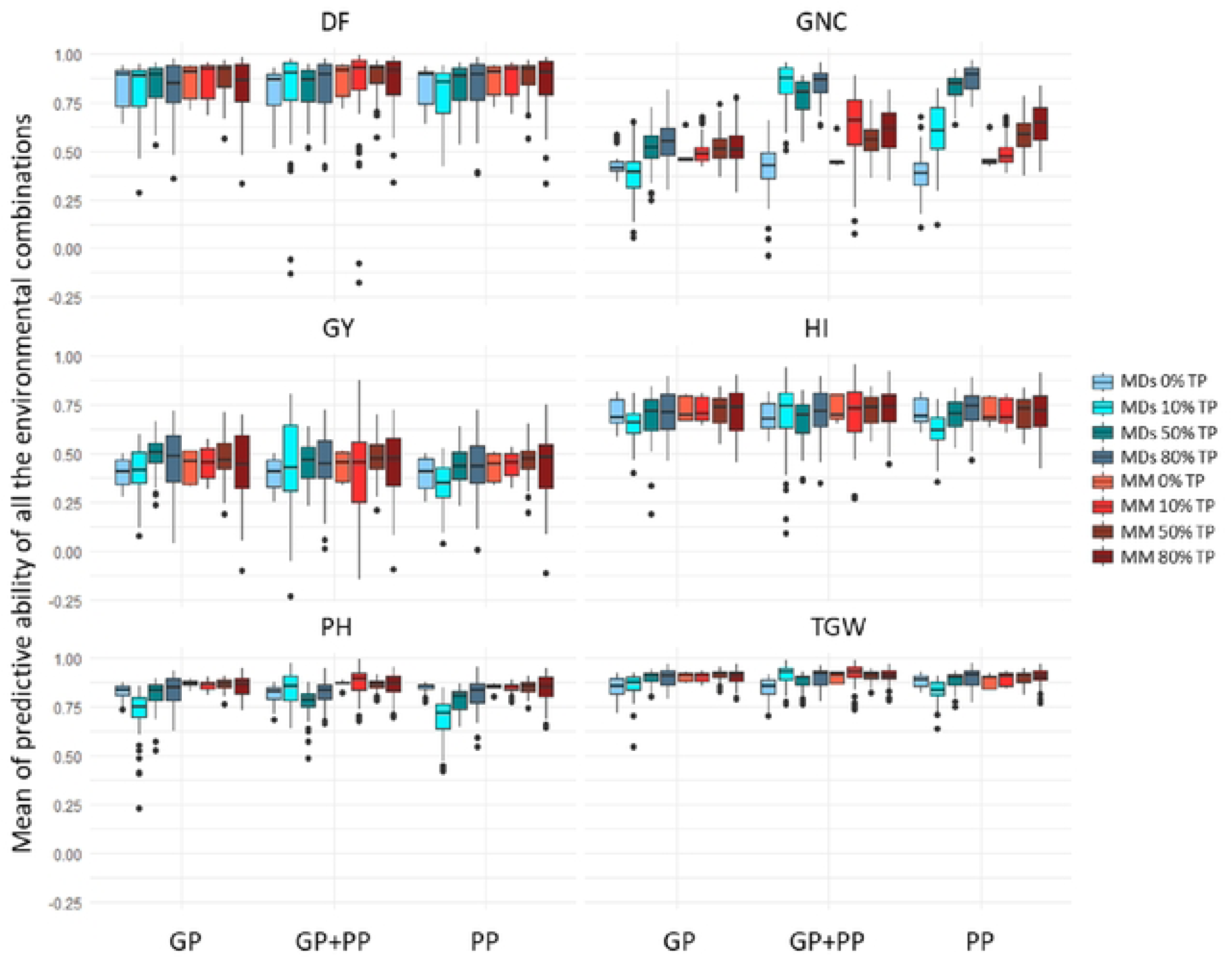
Means of the predictive ability for all the combinations of environments when two environments were considered in the training population. The blue boxes represent the MDs model, integrating a GxE effect, and the red boxes represent the MM model without GxE effect. For each model (MDs or MM) the difference of coloration of the boxes represents the percentage of genotypes of the second environment included in the training population (0%. 10%. 50% or 80%). the rest of the second environment being included in the validation population. The PAs arc averaged over rhe 6 scenarios tested; scenario 5 to scenario 10 (sec Table 1 for a detailed description of scenarios). The three prediction methods arc GP. genomic predictions; PP. phcnomic predictions; GP+PP. combination of genomic and phcnomic information for the predictions. DF. Day to flowering; PH. plant height; TGW. thousand grain weight; GY. grain yield; HI. harvest index; GNC. grain nitrogen content

Considering the MDs model with 50% of genotypes from the target environment in the TP (i.e., the 2-fold CV, scenarios 5 to 10), an assessment of the impact of the environment combinations was performed (Fig 4). No global trend emerged across all traits. PA were not increased when both environments were from the same year or from the same nitrogen treatment. Including a stressed environment in the TP, such as LN, did not always lead to increased PA in GP, PP or GP+PP.

**Fig 4.**
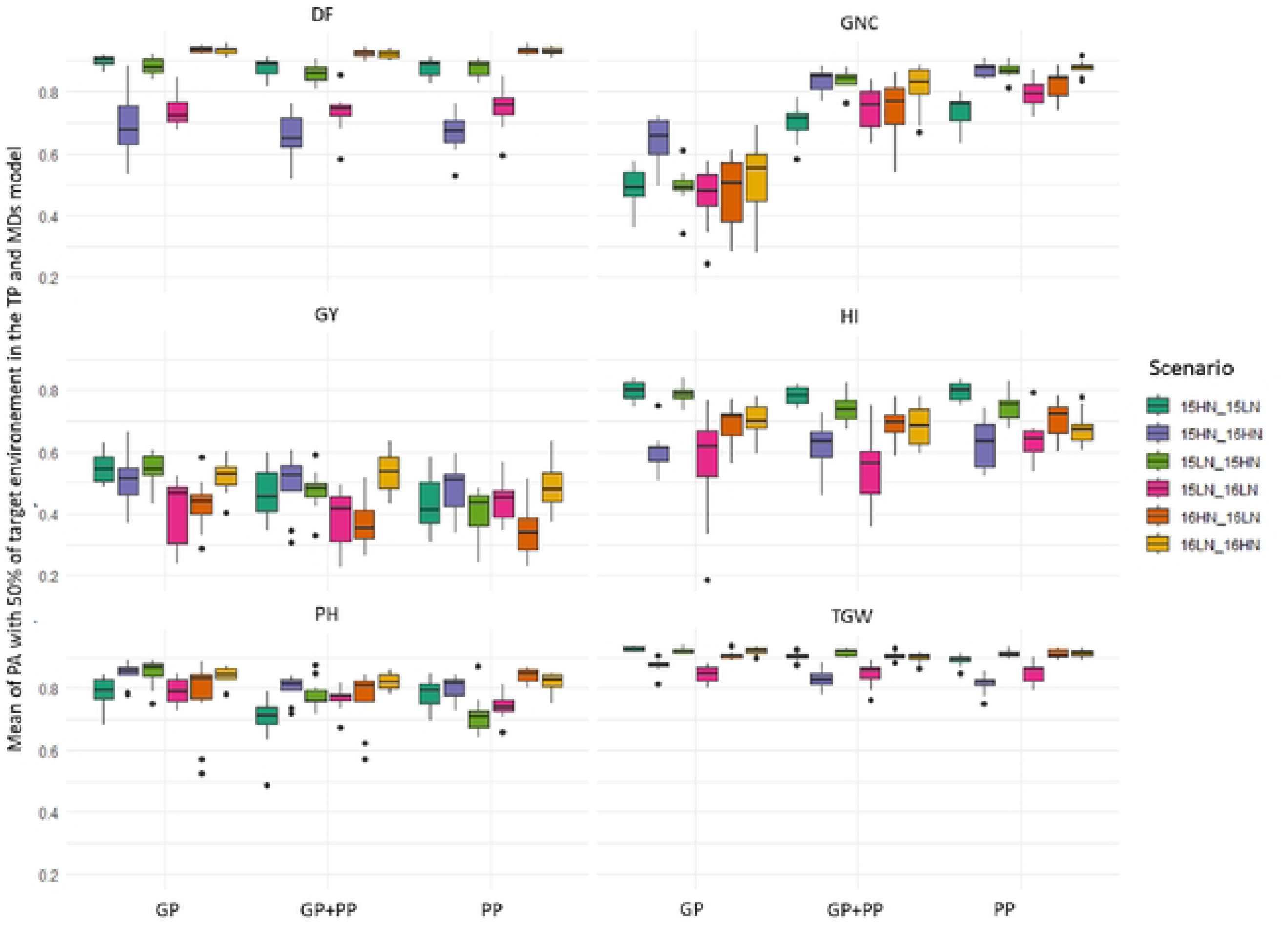
Means of the predictive ability achieved in the MDs model trained with a training population including 50% of the target environment (scenario 5 to 10) The three prediction methods arc GP. genomic predictions; PP. phcnomic predictions; GP+PP. combination of genomic and phcnomic information for the predictions. A detailed description of the scenario is given in Table 1. DF. Day to flowering; PH. plant height; TGW. thousand grain weight; GY. grain yield; HI. harvest index; GNC. grain nitrogen content

Minimal differences in PA were observed among GP, PP, and GP+PP for most traits, as illustrated where MDs model was used with a 2-fold CV in Fig 4. The exception was for GNC, where PP (0.609 ± 0.004) and GP+PP (0.640 ± 0.004) displayed significantly higher PA compared to GP (0.489 ± 0.004). For the other traits, even when statistically significant effects (p<0.05) were detected, the magnitude of the differences in PA between the best and worst performing methods (GP, PP, or GP+PP) remained low, below 0.025. For instance, for GY, the average PA for GP, GP+PP, and PP were 0.444 ± 0.005, 0.438 ± 0.005, and 0.422 ± 0.005, respectively. These minor variations highlight the comparable performance of all prediction methods based on genomic, phenomic or combined genomic and phenomic information for GY.

Using all 14 scenarios, the impact of relationship matrices (prediction models with GP, PP and GP+PP) and TP (one, two, or three environments integrated) was assessed, prioritizing these two parameters over the number of k-fold CV (here fixed at k=5 folds) or the type of model analyzed (here we chose the MDs, Fig 5). Whichever the relationship matrix used, for all traits except GNC, an increase in predictive ability (PA) was observed when the number of environments integrated in the TP increased from one to two. However, incorporating a third environment resulted in a plateau, with no further improvement in PA. The increase in PA observed when moving from one to two environments included in the TP, which corresponds to a shift from classical CV to sparse-testing validation, was greater for PP compared to GP and GP+PP (Fig 5). Yet, PAs eventually reached the same level for all three prediction models. For GNC, GP consistently gave lower PA than any combination involving PP, regardless of the number of environments included in the TP. However, for this trait, models with GP also benefit from a steady increase in PA (PA ranging from 0.4 to 0.6) with an increased number of environments which was not the case for the PP and GP+PP models.

**Fig 5.**
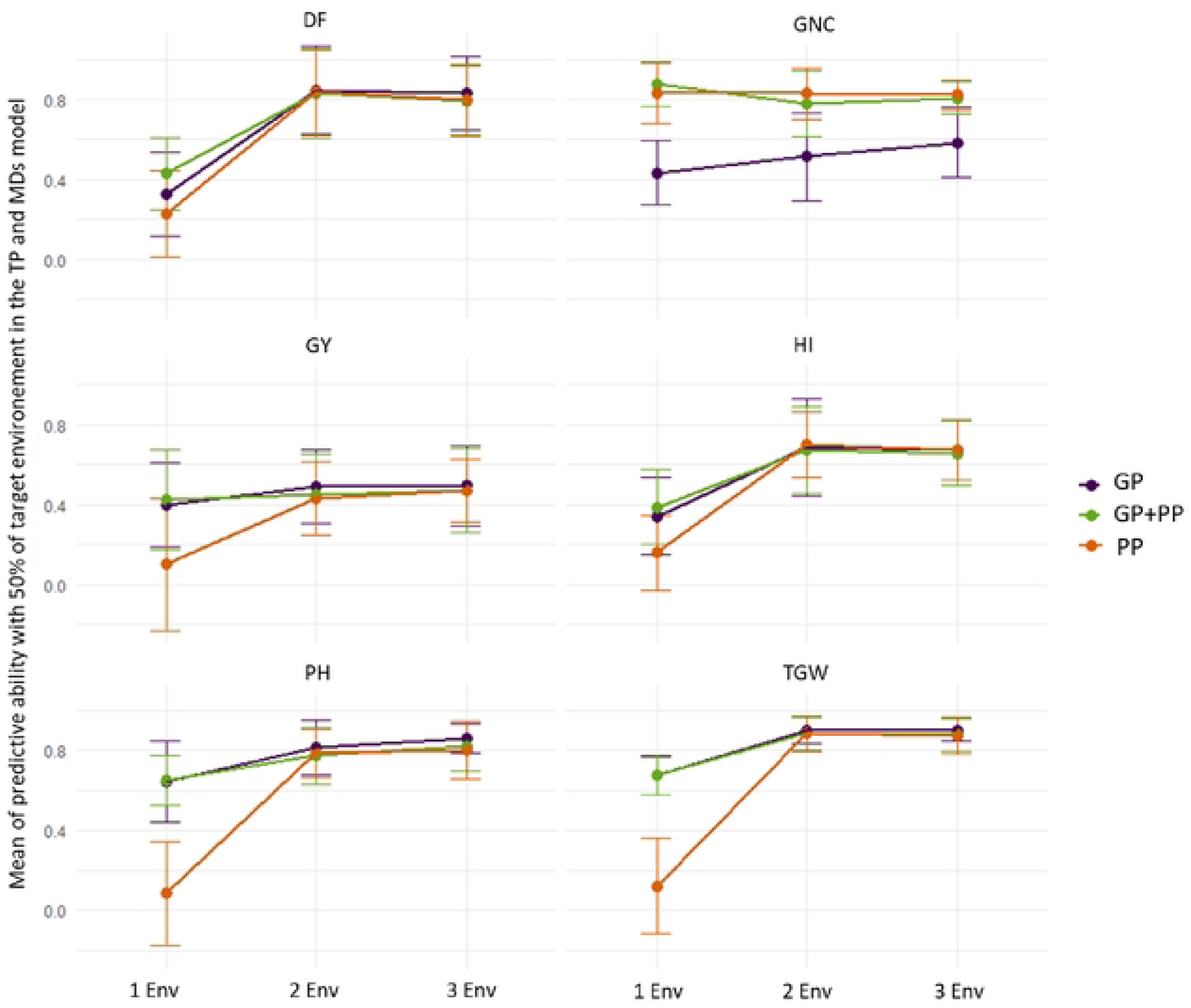
Means of the predictive ability for a training population including one. two or three environments, including 50% of the target environment and a MDs model, considering the three prediction methods (GP. genomic predictions. PP. phenomic predictions; GP+PP. combination of genomic and phenomic information for the predictions). The PAs arc averaged over all the scenarios tested within the corresponding group of scenarios i.e., scenario 1 to 4; scenario 5 to 10 and scenario 11 to 14 for Env I. Env2. and Env3. respectively (sec Table 1 for a detailed description of scenarios). DP. Day to flowering; PH. plant height; TGW. thousand grain weight; GY. grain yield; Hl. harvest index; GNC. grain nitrogen content

## Discussion

### Phenomic and genomic predictions comparison

Phenomic selection, a recent methodology introduced by Rincent et al. (10), shares a lot of similarities with GS. The distinction lies in the use of NIRS data, instead of single nucleotide polymorphism (SNP) data, to capture genetic information and assess genotype similarity. The effectiveness of PS has been demonstrated in diverse species and across various traits, with promising results reported in several studies (4,29). We compared the predictive ability (PA) of models including different relationship matrices with GP (genomic prediction using the genomic relationship matrix (GRM)), PP (phenomic prediction using hyperspectral relationship matrix (HRM)) and GP+PP the combination of both matrices in the prediction model. We thus compared the PAs of PP with GP and the combined GP+PP approach, explored the influence of training population size and composition and investigated the impact of incorporating GxE effects into different models. To our knowledge, the present study is the first to evaluate the applicability of PS in rice, despite the importance of this species worldwide.

When analyzing single environment models with conventional k-fold CV, GP were slightly higher than what was previously estimated in the rice literature for DF, PH and GY (45–47). Phenomic prediction exhibited substantially lower PA compared to GP or combined GP+PP, with the exception of GNC for which NIRS based calibrations are used as a proxy of the trait (48). This result deviates from the literature, where PA using HRM and GRM are often comparable across various traits and species (10,29,42). To address potential concerns regarding the tissue or NIRS measurements, we conducted pre-analyses on milled dry straw (data not shown) that yielded similar results to the analyses using milled dry grains. This suggests that the observed low PA based on PP in single-environment were not due to tissue-specific effects or NIRS measurement issues. Other pre-analyses were performed with different NIRS data preprocessing (data not shown) as it was previously highlighted that PP could be influenced by the preprocessing applied on the spectra (42). However, none of the pre-processing methods yielded significant improvement in PA for PP compared to GP. The most plausible explanation for the low PA in PP likely stems from the generally low heritability associated with the NIRS wavelengths employed in this study (S1 Fig.). The NIRS data may not have captured a sufficient amount of genetic information relevant to the target traits compared to SNP data. Spectra with low wavelength heritability gave generally lower PA than spectra with high heritability (17).

### Factors influencing phenomic preditions: effects of the training population size and multi-environment models

The present study also examined the influence of TP size on PA. As expected, increasing TP size led to improved PA, consistent with previous observations by Galan et al. (33). However, the magnitude of improvement was not substantial, as observed by Robert et al. on wheat (44). These authors demonstrated a marginal increase PA for grain yield, ranging from approximately 0.5 with a TP size of 50 to around 0.6 with a TP size of 250. No real differences in the effect of TP size were observed between GP, PP, or the combined GP+PP approach in the present study. While several studies (30,49) reported that PP requires a smaller TP size for comparable PA compared to GP, Zhu et al. (30) attributed this to the presence of multiple values per wavelength in NIRS data, as opposed to the binary nature of SNPs in genomic data. Unlike marker data which provides discrete genotypic categories (two genotypic classes or three if heterozygotes are present) at each marker, NIRS data offers a continuous distribution of reflectance values for each wavelength, representing a wider range of potential states. This suggests that NIRS data might enable achieving similar PA with fewer genotypes in a breeding program, potentially reducing phenotyping efforts, although this was not found with the present results. This could be explained by the lower number of genotypes included in the analyses in the present study compared to the one of Zhu et al (30).

Interestingly, incorporating data from an additional environment into the TP significantly improved PA for PP, bringing it at the same level of prediction as GP or GP+PP, which was in accordance with the literature (17,32). The inclusion of NIRS data from additional environments within the models likely enhances the estimation of genotype relationships, as previously demonstrated by Robert et al. (32). This improvement may be attributed to the high correlations observed for the phenotypic traits between the environments and the relatively high heritability of those target traits. In essence, the strong environmental correlations may have allowed PP, in this specific case, to achieve a level of PA comparable to GP. Zhu et al. (30) reported that for triticale, the PA of PP was highest when environments exhibited strong correlations. This observation is further supported by the work of Robert et al. (32), who investigated PP in wheat considering four datasets in different MET scenarios. Their findings suggest that the improvement in PA achieved by incorporating multiple environments, compared to a single one, depends on the degree of correlation between those environments for the trait of interest. This dependence likely arises because the HRM is likely to be more comparable between correlated environments than between uncorrelated ones. Nevertheless, it is important to acknowledge that even with lower heritability or weaker correlations between environments, PP can offer valuable applications. Indeed, as highlighted by Lane et al. (29), PP can be effective for ranking genotypes relative to each other. Robert et al. (32) further highlighted that genotypes can exhibit distinct spectral profiles in one environment but similar profiles in another environment. This shows the potential of incorporating hyperspectral data from multiple environments to enhance PA in PP. These variations in genotype specific spectra across environments likely reflect the capture of some GxE interaction effects. This explanation aligns with the observation in the present study that models incorporating GxE effects (MDs model) yielded similar or even superior PA than models without GxE effects for four of the six traits measured in the present study. Unlike molecular markers, NIRS data are susceptible to environmental variations. The potential benefits of incorporating GxE effects into GP have also been emphasized by several studies (17,19,22,27,32,50). Our study reinforces this result and extends it to PP. The observed improvement in PA with GxE models in the present study could also align with the hypothesis proposed by Robert et al. (32) that variance of the traits were not only additive but also interactive (GxE).

The inclusion of a third environment into the TP did not lead to a continuous increase in PA. Instead, PA appeared to plateau after incorporating two environments. This observation suggested a redundancy effect associated with including excessive environmental data (32). In the context of breeding programs, it is crucial to strike a balance between the cost associated with NIRS measurements in multiple environments and the potential gains in PA. Future research efforts could explore cost-effective strategies to optimize the number of environments used in PP for breeding purposes.

### Influence of combined information use

It is important to note that the models employed in the present study differed from those proposed by Robert et al. (32) or Lane et al. (29) in their treatment of GxE effects and multi-environment hyperspectral data. In the first study, NIRS datasets were combined at the analysis outset, leading to the estimation of a single HRM that integrated information from all environments, similar to a GRM. Lane et al. (29) adopted a contrasting approach, estimating a separate HRM for each environment and subsequently averaging them to create a single matrix for prediction. In contrast, the present study aimed to capture the genotypic effects associated with each environment. To achieve this, the number of genotypic effects was set equal to the number of environments, with one random effect added for each NIRS dataset. This methodological choice likely influences the predictions by capturing the variance explained by genotypic effects differently compared to the aforementioned studies. The MDs model employed in the present study represents a block diagonal structure, where distinct matrices are utilized for each environment with a similar GxE variance between the environments (16). This approach does not allow to take advantage explicitly from the HRM of correlations between environments. Consequently, the model did not capture or integrate the covariance matrices that might exist between environments. Investigating the PA of alternative models that incorporate these variance-covariance matrices between environments could be a promising avenue for future research (51,52). Such models might offer the potential for enhancing even more the models’ PA.

Present study investigated the potential benefits of combining GP and PP by incorporating them as separate genotypic random effects into the models. While this approach improved PA, the increase was not substantial across traits, number of environments, or TP size. Overall, the combined GP+PP strategy resulted in only a slight improvement in PA compared to any of the best performing method among GP and PP models. These findings align with observations by Robert et al. (32), who reported also only slightly enhanced PA compared to genomic models when combining molecular and hyperspectral data in their models. However, unlike the latter study which employed a single combined HRM-GRM relationship matrix, the present study utilized separate matrices. In addition, it is also important to stress that despite the observed improvement in PA, its magnitude may not justify the additional cost associated with combining both methods in a breeding program. Therefore, the optimal choice between GP and PP might depend on the specific traits of interest considering cost-effectiveness.

Previous research suggests that incorporating NIRS data collected under stress or unfavorable conditions can enhance PA for PP (10,29). In the context of the present study, although the low levels of nitrogen fertilizers might be considered analogous to such stress conditions, they did not provide improvement of the PA when considered in the TP. Interestingly, Lane et al. (29) observed that the water-stressed environment, which yielded the highest PA in their study, also exhibited the lowest repeatability. They mentioned that optimal TP composition for PP would encompass a broad range of genetic diversity and data from multiple years. This conclusion aligns with the notion that genotypes tend to exhibit greater phenotypic variability in stressed environments compared to non-stressed conditions. To evaluate the added value of incorporating NIRS data from stressed environments into the TS, it is important to assess performances of phenomic predictions across a broad spectrum of environments. This assessment should encompass a large panel of environments with a wide range of correlations between them. Such an investigation would provide a more robust understanding of the generalizability and effectiveness of this approach and would help breeders to be more efficient in the predictions when using phenomic selection.

### Breeding program applications

Present study represents the first investigation comparing the efficiency of PP to GP in rice. A significant advantage of PP over GP lies in its reduced infrastructure requirements (53). Notably, PP only necessitates a NIRS spectrometer, which can lead to substantial cost reductions. While NIRS material can be quite expensive, it is frequently already available in the centers or institutions running breeding programs. Van Tassel et al. (54) estimated genotyping costs to range from $30 to $50 per genotype, compared to $2 to $15 per genotype for NIRS measurements, depending on the technology employed. Even if two NIRS measurements of each genotype are needed to achieve comparable accuracy to genomic selection, costs would still be between $4 to $30 per genotype (to which would have to be added the costs of producing the genotypes in a second environment), making NIRS a more cost-effective approach. Furthermore, PP has the potential to increase the accuracy of GxE effect estimation, potentially leading to a more accurate evaluation of genotype value. Additionally, NIRS is a non-destructive method and measurements can be performed on grains prior to sowing, which can contribute to increased selection intensity.

While PP offers advantages, it also raises several questions not encountered with GP. A critical consideration involves the optimal tissue for NIRS measurement. Leaf-based measurements are appealing due to their ease and feasibility in field settings without harming plants or requiring harvest. This approach could facilitate early selection before harvest or even flowering (allowing to identify the most relevant crosses to perform in recurrent selection approaches), potentially reducing phenotyping costs and breeding cycle length. As NIRS can be affected by environment, which makes it a good option to capture GxE, would it also be sensitive to physiological stages when the data is collected?

Furthermore, previous research suggests that for PP, tissues more closely related to the target trait tend to yield higher PA (53). Supporting this notion, Robert et al. (44) reported lower prediction accuracy for GY using leaf spectra compared to grain spectra.

Another important question to consider is the optimal integration point of PP within the breeding scheme compared to GP. Rincent et al. (10) suggested that PP holds promise for selecting superior genotypes prior to multi-environment testing. In this context, predicting phenotypic values might offer advantages over predicting additive genetic values. Future research employing simulations could address this specific question by evaluating the impact of key factors known to impact the efficiency of breeding schemes. These factors could include the number of genotypes under study, the number of environments considered, and the optimal timing of PP implementation relative to GP within the breeding cycle.

## Material and Methods

Details concerning the experimental population, field experiment design, hyperspectral measurements, and genotyping procedures can be found in Rakotoson et al. (43,55) and Rakotomalala et al. (56). A summary of these aspects is provided in the following sections.

### Experimental population

The experimental population consisted of a panel of 190 rice accessions that represents the working collection of the upland rice breeding program conducted in Madagascar by FOFIFA (Malagasy National Research Center for Rural Development) and CIRAD (French Agricultural Research Center for International Development). The majority of these accessions came from the tropical japonica genetic group and originated from various breeding programs, including the FOFIFA-Cirad program, or historical acquisition from IRRI (International Rice Research Institute), CIAT (International Center for Tropical Agriculture in Colombia), EMBRAPA (Empresa Brasileira de Pesquisa Agropecuária in Brazil) and Africa Rice (S4 Table).

### Field experiments

Field trials were conducted over two cropping seasons (2014-2015 and 2015-2016), hereafter referred to as 2015 and 2016, at Ivory in Madagascar (19°33’27” S, 46°24’43” E, 960 masl). A split-block design with two replications was employed in a randomized alpha lattice for each year. Each replication consisted of 14 blocks, further subdivided into two sub-blocks for evaluating nitrogen fertilization levels. Within each block, 16 accession plots were sown, including 14 tested and two control accessions. Each accession plot received 5 tons of manure per ha and was further divided into two adjacent subplots, one receiving no mineral nitrogen fertilization (LN for low nitrogen) and the other receiving additional mineral nitrogen (HN for high nitrogen). Subplot sizes varied slightly across replications and years. In 2015, subplots measured 1.8 m × 2.4 m in the first replication and 1.4 m × 2.0 m in the second replication. In 2016, subplots were 1.8 m × 1.6 m in the first replication and 1.2 m × 1.6 m in the second replication. Field preparation began with traditional ox plowing, followed by hand-leveling of the soil surface. Sowing involved sowing four to six rice seeds per hill at 20 cm intervals in both directions. A standardized base dressing was applied to all plots before sowing, incorporating cattle manure (5000 kg.ha^-1^), triple superphosphate (69 kg.ha^-1^ P2O5), potassium sulfate (62.4 kg.ha^-1^ K2O), and dolomite (500 kg.ha^-1^). For the HN plots, additional nitrogen fertilization was provided using urea (46% N) at a total amount of 120 kg.ha^-1^, split into three equal applications at the emergence, tillering, and booting stages. Phytosanitary treatments were applied as required for growing rice in the area and to protect the crop from pest and diseases. As the data included in this study were collected across two years (2015 and 2016) and two nitrogen treatments, the four nitrogen-year combinations were henceforth referred to as environments.

### Phenotypic trait measurements and phenomic data

Building upon the work of Rakotoson et al. (43), this study focuses on six key traits previously evaluated in each elementary plot. These traits were chosen for their relevance to breeding programs and their diverse genetic architectures. Days to flowering (DF) was recorded when 50% of the plants within the plots reached flowering (in days). At maturity, plant height (PH) was measured on six plants located in the middle of each plot averaged and expressed in cm. Then, the panicles were counted and hand-threshed, and filled grains were separated from unfilled grains. The dry weight of filled grains was determined after oven-drying at 60°C for 72 h. Filled grains were used to estimate grain yield (GY) calculated and expressed in kg.ha^-1^. Two sub-samples of 200 filled grains were used to calculate thousand-grain weight (TGW) expressed in g. Harvest Index (HI) was calculated as the ratio between GY and total dry biomass (straw and grain yield). Finally, grain nitrogen content (GNC; %) was measured by collection of near-infrared spectroscopy (NIRS) on dry grains using a monochromator (LabSpec 4 Standard-Res Lab Analyzer, ASD Inc., Boulder, USA; wavelengths 1 000–2 500 nm). These samples were previously ground to 1mm, using a model 1093 Cyclotech sample mill (FOSS, Höganäs, Sweden). Near infrared spectra were acquired on 4 samples (technical replicates) per grain lot and the average spectra were calculated. The NIR spectra of each accession included data from 1,500 wavelengths between 1,000 and 2,500 nm with a 1 nm step.

### Genotypic data

Seeds from each accession were grown for DNA extraction at CIRAD Montpellier laboratory in France. Leaf tissue from a single plant per accession was used for DNA extraction using the MATAB method (57). The extracted DNA was then diluted to a concentration of 100 ng.μL^-1^. Each DNA sample was digested with the ApekI restriction enzyme. Then, each library was single-end sequenced in a single-flow cell channel (i.e., 96-plex sequencing, (58)) using an Illumina, Inc. HiSeqTM 2000. Reads were aligned to the rice reference genome (Os-Nipponbare-Reference-IRGSP-1.0, (59) with Bowtie2 (default parameters). SNP calling was performed using the Tassel GBS pipeline v5.2.37 (default parameters, (60)). SNPs, with a call rate < 80%, a heterozygosity rate > 20% or a minor allele frequency (MAF) < 2.5%, were all discarded. The remaining heterozygotes were converted into missing data. Then, missing data were imputed using Beagle v4.0 (61). After imputation, markers with a MAF < 4.2% (8 out of 190) were discarded. The final resulting matrix comprised 190 individuals and 38 079 SNP markers.

### Statistical analyses

#### Phenotypic data analyses

The raw data were checked per environment for outliers using the boxplot.stats function of the R package “stats” (62) with a coefficient of 1.5, which means that outliers were identified if the phenotypic values were outside 1.5 times the interquartile range above the upper quartile and below the lower quartile. No outliers were discarded. The following mixed model was used for variance decomposition, as it was the most parsimonious according to the AIC criterion:

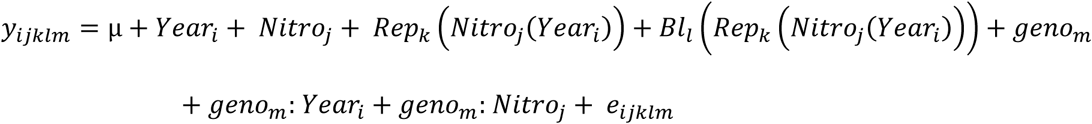

Where *y*_*ijklm*_is the vector of phenotypic values, µ is the overall mean of the phenotypic value, *year*_*i*_ is the fixed effect of the year (2015 or 2016); *Nitro*_*j*_ is the fixed effect of the nitrogen treatment (HN or LN); *Rep*_*k*_(*Nitro*_*j*_(*year*_*i*_) is the fixed effect of the replicate within a year and a nitrogen treatment; *Bl*_*l*_ (*Rep*_*k*_(*Nitro*_*j*_(*year*_*i*_)) is the random effect of the block l nested in replicate k and nitrogen treatment j and year i with distribution *Bl*∼*N*(0, σ^2^_g_); *geno*_*m*_ is the random effect of the genotype m with distribution *geno*_*m*_∼*N*(0, σ^2^_Bl_); *geno*: *year* is the random effect of the genotype m by year i, which is part of the genetic by environment interaction effect; *geno*_*m*_: *Nitro*_*j*_ is the random effect of the genotype m by nitrogen treatment j, which is the second part of the genetic by environment interaction effect; and *e*_*ijklm*_ is the residual considered as a random effect with distribution *e*∼*N*(0, σ^2^_e_). Variance decomposition was performed using the lmer function of the R package “lme4” (63).

Broad sense heritability (H^2^) was estimated using the following equation:

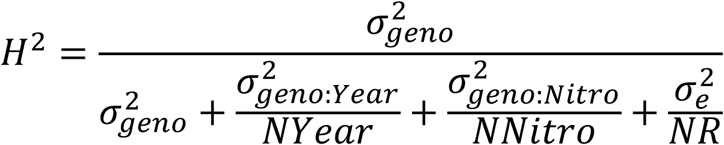

where σ^2^*_geno_* is the variance associated with genotypes, σ^2^*_geno:year_* is the genetic by year interaction effect variance, σ^2^*_geno:nitro_* is the genetic by nitrogen treatment interaction effect variance, σ^2^ is the residual variance, NYear and NNitro are the harmonic mean of the number of years and nitrogen treatments per genotype and NR is the harmonic mean of the number of replicates per genotype across the years and nitrogen treatments.

To estimate the correlations between environments, for each trait, correlations of phenotypic values between the two years and two nitrogen treatments (four environments) were performed using the rcorr function of the R package “Hmisc” (64).

#### Spectra pre-processing

As detailed by Brault et al. (37), spectra were processed separately within each environment. From the average spectra, several pre-processing methods were performed (e.g. smooth, standard normal variate or normalization, detrend and first and second derivative on normalized spectra). From the preliminary analyses, it appeared that pre-processing with the second derivative spectra was more relevant than the other pre-processing methods to perform predictions. Consequently, we only kept this pre-treatment for the rest of the study. A mixed model over the reflectance at each wavelength was computed to estimate the variance components and derive NIRS genotypic BLUPs. The mixed model equation was:

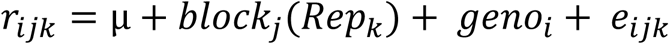

With *r*_*ijk*_ the reflectance at a given wavelength, µ the intercept; μ*block*_*j*_(*Rep*_*k*_) the random effect of block j nested in the repetition k; *geno*_*i*_ the random genotypic effect of genotype i, and *e*_*ijk*_ the residual. NIRS heritability (H²) for each wavelength was estimated using the following equation:

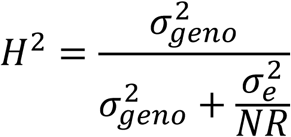

where σ^2^ is the variance associated with genotypes and NR is the harmonic mean of the number of replicates per genotype.

#### Genomic and phenomic predictions

Genomic prediction models were developed based on a two-stage procedure. First, to correct for the random effects of the replicates and blocks (*Bl*_*j*_(*Rep*_*k*_)), best linear unbiased estimations (BLUEs) of the genotype effect were estimated for each trait within each environment (combination of year and treatment) using the lmer function (from the lme4 package) and the following model:

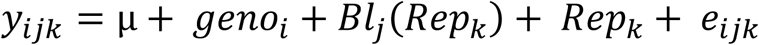

Where *geno*_*i*_ is the fixed effect of the genotype i.

The GP models were run by year and nitrogen treatment and the BLUE values for each trait were used to compare with the predictions.

To assess the relative effectiveness of PP compared to GP, various cross-validation scenarios were tested. These scenarios explored the influence of training population (TP) and validation population (VP) composition on the predictive accuracies (PA), as detailed in Table 3. In addition, the scenarios varied based on the number of environments included in the analyses.

When a single environment was considered, we implemented a k-fold cross-validation approach (65) with k sets to 5 (80% of the population used for training and 20% for validation) or k set to 2 (equal split of the population for training and validation). For each case, 10 replications were performed, where all the genotypes were predicted using the 5- or 2-folds cross validation and correlations between predicted values and BLUEs were estimated for all the genotypes together.

For scenarios involving two environments, data from the first environment were always incorporated entirely into the TP. Additionally, a variable portion of data from the second environment was also included in the TP to assess the impact of the training set size. This variable portion ranged from 0% (no information from the second environment) to 10%, 50%, and 80%. These variations allowed to evaluate the combined influences of the TP size and of the inclusion of information from an additional environment and also to compare these scenarios to those using only one environment.

Scenarios involving three environments followed a similar structure to those with two environments. However, data from the second environment was entirely incorporated into the TP in all three-environment scenarios. The variable portion for the third environment included in the TP ranged from 0%, 50%, and 80%. The remaining data from the third environment (100%, 50%, or 20%, respectively) constituted the VP All the details of the environment combinations are provided in Table 3.

Bayesian linear mixed model was performed for all the analyses using the R package BGGE (66). For scenarios including one environment, the genomic prediction (GP) and phenomic prediction (PP) were run using a univariate single-environment model (SM) G-BLUP model considering only the main genotypic effects.

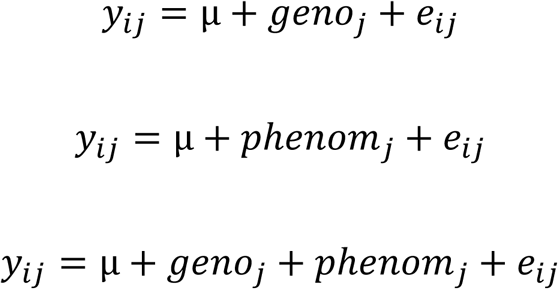

Where *y*_*ij*_ is the vector of genotype BLUEs, µ is the intercept and *geno*_*j*_ a vector of random genotypic effect normally distributed *geno*_*j*_∼*N*(0, σ^2^*g*), with K a kinship matrix based on SNP marker information calculated with the formula proposed by VanRaden (67) for GP, and *phenomm*_*j*_ a vector of random phenomic effect normally distributed *phenomm*_*j*_∼*N*(0, σ^2^_g_*H*) with *H* a hyperspectral relationship matrix defined as such *H* = *S_p_*^∗^ ∗ *S_p_*^∗′^⁄*L* where *S_p_*^∗^ is the matrix of NIRS genotypic BLUPs and L the number of wavelengths, for PP. The third model combine genomic and phenomic information.

In the scenarios including more than one environment, a fixed effect of the environment (MM model) or a G×E interaction random effect (MDs model) was added to the predictive model. To do so, G×E genomic variance matrices were constructed.

The multi-environment model (MM) assumes that genetic effects across the environment are constant across genotypes, and therefore the absence of G×E. In this model, a single matrix containing the genomic relationships was constructed for the main across-environment effects:

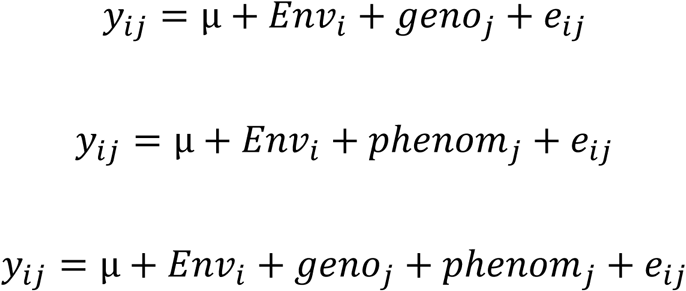

with *Env*_*i*_ the fixed effect of the environment (year x nitrogen treatment interaction), *geno*_*j*_ the random effect of the genotype, having a variance-covariance structure following *G*∼*N*(0, *J*⊕*g*σ^2^), where *J* is a Nenv x Nenv unit matrix (full of ones) and *K* similarly defined as before. The Kronecker product between the unit and the relatedness matrix (*J*⊕*g*) can be interpreted as a uniform genetic effect across environments. *phenomm*_*j*_∼*N*(0, *J*⊕*H*σ^2^) is defined in a similar with *H* a hyperspectral relationship matrix representing the phenomic information.

The multi-environment model (MDs) which is an extension of the MM model includes a single random deviation effect of the G×E:

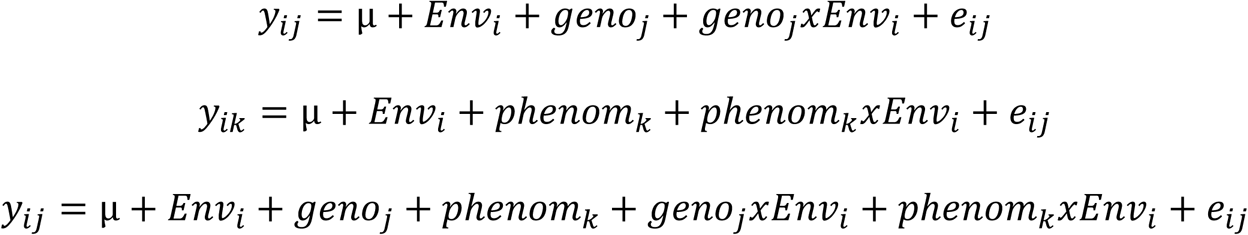

With all terms similarly defined as in the previous models and *geno*_*j*_(*Krphenomm*_*k*_)*NEnv*_*i*_∼*N*(0, σ_*Nenewv*_ ⊕*g*(*H*)σ^2^) representing a uniform deviation due to environmental influence, for the genomic (K) or phenomic (H) terms respectively.

Full details about these models can be found in Granato et al. (66). All genomic predictions were performed using the R package BGGE (66) with the following parameters: burn-in = 10,000, nIter = 70,000 and thin = 10.

Predictive ability (PA) for each model (GP, PP or GP+PP), scenario and TP size was assessed by calculating the correlation between predicted values and the BLUEs. To account for potential stochastic effects influencing accuracy comparisons between models and scenarios, all predictions were replicated 10 times considering different training populations. This allowed to estimate and compare the mean and standard deviation of PA for each model across all scenarios.

Linear mixed models were then used to compare the model predictive ability. These models considered the fixed effect of the matrix, training population size, model, or environments or combination of effects according to the analyses and included a Z-transformation of the PA data to ensure normality.

## Acknowledgements

The authors would like to thank the excellent FOFIFA field staff in Madagascar. We would also like to thank all the persons that participated in data acquisition and sharing during this GS-Ruse project, jointly supported by Cariplo (Italia) and Agropolis (France) Foundations (Grant No. 1201-006).

## Supporting information

S1 Table: Descriptive statistics of the phenotypic measurements performed in each of the four environments

S2 Table. Results of the LMM analyses comparing all the effects for all the scenarios and all the traits, using Z-transformed PA.

S3 Table. Predictive ability of the different scenarios and models (means ± standard deviation).

S4 Table. List of the rice accessions used

S1 Fig. Wavelength heritability after der2 pre-processing within each environment

## Notes

### Competing Interest Statement

The authors have declared no competing interest.

